# The gut microbiota is a determinant of sexual dimorphism in ALS-linked TDP43 mice

**DOI:** 10.1101/2024.08.29.610355

**Authors:** Evandro J. Beraldi, Sukyoung Lee, Yulan Jiang, Isabel M. Rea, Rushda Phull, Catherine M. Keenan, Matthew Stephens, Martin Bardhi, Oluwamolakun Bankole, Gerald Pfeffer, Jeff Biernaskie, Kathy D. McCoy, Eran Elinav, Keith A. Sharkey, Minh Dang Nguyen

## Abstract

The mechanisms underlying the earlier onset and male predominance of amyotrophic lateral sclerosis (ALS), the most common form of human motoneuron disease, are poorly understood. Here we show that the gut microbiota protects against TDP43 toxicity and contributes to the sexual dimorphism in mice expressing a mutant form of TDP43 (A315T) linked to ALS. TDP43 mice raised under germ-free conditions, or treated with antibiotics to deplete the gut microbiota, develop motoneuron disease earlier and show no sex differences in onset and lifespan. Behavioral and histopathological analyses confirm the exacerbation in neurodegeneration caused by the absence of gut microbiota. Castration did not alter disease course of male TDP43 mice, suggesting that male sex hormones do not interact with the gut microbiota to confer disease phenotype. Future identification of gut bacteria species and their mechanisms of action offers a unique opportunity to understand sexual dimorphism in ALS, with the ultimate goal to develop non-invasive and sex-specific treatments for ALS.

## Introduction

Amyotrophic lateral sclerosis (ALS) is the most common form of human motoneuron disease; it is caused by the degeneration of upper and lower motor neurons, resulting in progressive weakness of voluntary muscles, and early death, typically about 3 years from disease onset ^1–6^.The lifetime risk to develop ALS is around 0.3% ^7^. Males are up to two times more susceptible than females to develop ALS ^2^, but this propensity varies as a function of age, lifestyle and geographical location. Furthermore, the age of onset is also earlier for males than females with ALS ^8^, making ALS the most rapidly progressive neurodegenerative disorder with sexually dimorphic features. The mechanisms underlying sexual dimorphism in ALS are not well understood (reviewed in ^9^), with hormones being commonly viewed as the main driver for sex differences ^9^.

Recent evidence suggests that the gut microbiome (the ensemble of microorganisms including bacteria, viruses, protozoa, fungi and archaea that mediate the nutritional, metabolic and immune/host defense functions of the organ) ^10–12^ contributes to the pathogenesis of ALS in mice ^13–22^. A recent study from the Feldman group also reveals that that differences in gut microbiome composition between ALS and control patients are linked to plasma metabolites and particularly to lipids, with the potential to become a target for therapeutic interventions in human ALS ^23^. Work in a *C. elegans* model of ALS also demonstrated that *Lacticaseibacillus rhamnosus* HA-114 is neuroprotective through fatty acid metabolism and mitochondrial β-oxidation ^24^. In mice expressing a mutant form of SOD1 linked to familial ALS, depletion of gut bacteria exacerbates disease while re-establishing gut homeostasis with bacterial gut colonization or treatment of metabolites derived from protective bacteria slowed down disease, and even extended the lifespan of the mutant mice ^13,14^. The lifespan of C9ORF72 null mice, linked to ALS and fronto-temporal dementia (FTD), is also strongly dependent on the environment-controlled microbiota composition ^16^. Two strains of the same mouse line display strikingly different lifespans when housed in different facilities; fecal material transfer between the two strains reciprocally alters their survival ^16^. Thus, the environment and gut microbiota are complicit in ALS pathogenesis and the genetics of the host determines the impact of these variables on disease expression.

In this study, we address the uninterrogated but critical question as to whether the gut microbiota contributes to sexual dimorphism in ALS. This is an important topic because, in humans, the gut microbiota composition varies significantly between males and females ^25,26^, but the implication of this for disease expression and the identification of potential biomarkers and therapeutics in ALS remains ill defined.

We used transgenic mice expressing a mutant form of the TAR DNA binding protein 43 (TDP43 A315T) ^20,27–29^ found in ALS patients. The model has just begun to be studied in the context of microbiota ^20^, but importantly, has the potential to recapitulate key sex differences of the human disease ^2,8,9^. Nearly all human ALS cases (with or without TDP43 mutations) show the abnormal accumulation of TDP43, making this pathological signature a universal feature of both familial and sporadic forms of ALS ^30,31^. Using a combination of germ-free conditions and antibiotic treatment, behavioral, molecular and cellular approaches, we demonstrate a critical role for the gut microbiota in the pathogenesis and sexual dimorphism of TDP43 mice.

## Results

### Motoneuron degeneration occurs prior to gut obstruction in TDP43 A315T mice of both sexes

One of the key characteristics of ALS is its male predominance ^2,8,9^. This sexual dimorphism is recapitulated in TDP43 A315T mice, with males displaying accelerated motoneuron disease, and much shorter lifespan than females ^27–29^ . However, these studies have also documented severe gut obstruction and enlargement, leading to early lethality, particularly in males, precluding the development of motoneuron disease ^27–29^. We obtained TDP43 A315T (designated TDP43 herein) mice from Jackson laboratories, raised them in our specific-pathogen free (SPF) facility at the University of Calgary, and fed them with standard diet. In our mice, the mutant human protein is expressed in the brain, spinal cord and gastrointestinal (GI) tract using western blots (Figs. 1A, 2A). The mutant protein accumulates in cytosol and nucleus of spinal motoneurons (Fig. 1B), as well as in cholinesterase-positive neurons of the dorsal motor nucleus of the vagus nerve (not shown) and myenteric neurons in the gut (Fig. 2G), consistent with protein expression driven by a neuron-specific promoter.

**FIGURE 1.**
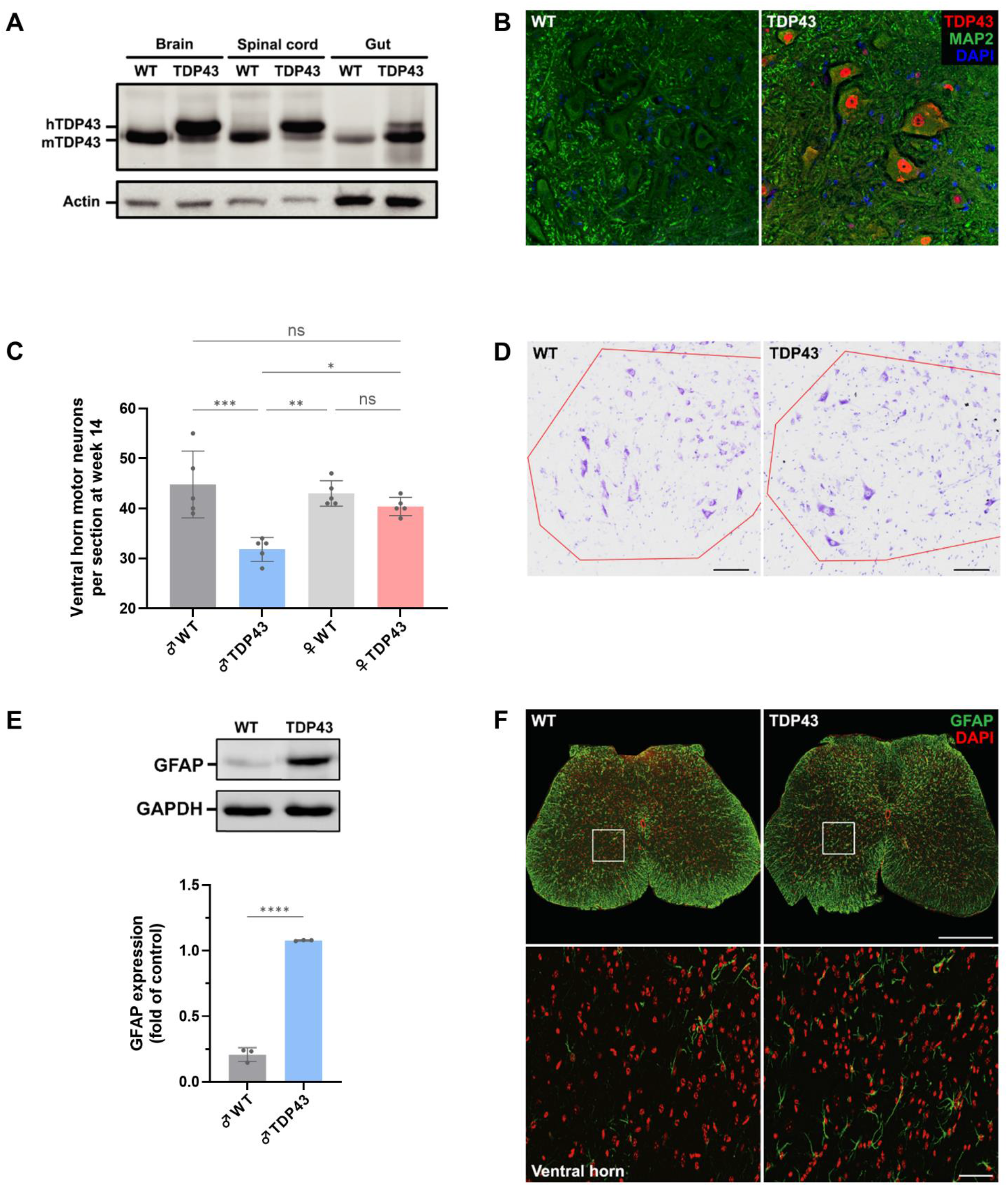
Motoneuron degeneration in ALS-linked TDP43 mice. (A) Representative western blot images showing expression of endogenous mouse TDP43 (mTDP43) and human mutant TDP43 (hTDP43) protein in wild type (WT) and TDP43 mice tissue (brain, spinal cord and gut lysates). (B) Confocal images of human mutant TDP43 protein (TDP43) immunofluorescence in the ventral horn of the lumbar spinal cord of WT and TDP43 mice. C) Nissl-stained motoneuron counts in the ventral horn of the lumbar spinal cord of 14 weeks old TDP43 mice of both sexes and age/sex-matched control littermates. Only TDP43 male mice show loss of large motoneurons at this age. Counts were made from 20 sections (16 μm) per animal, results are shown as mean ± SD, n = 5, \**p* < 0.05, \*\**p* < 0.01, \*\*\**p* < 0.001, one-way ANOVA test with posthoc Tukey’s test. (D) Representative images of Nissl-stained motoneurons in the ventral horn area (highlighted in red) of matching lumbar spinal cord sections of WT and TDP43 mice. Scale bar = 100 µm. (E) Representative western blot image showing glial fibrillary acidic protein (GFAP) protein expression in the spinal cord of WT and TDP43 mice at end-stage and quantification of the western blot. The intensity was normalized to GAPDH expression. Data are shown as mean ± SD, n = 3, \*\*\*\**p* < 0.0001, unpaired t-test. (F) Confocal images of GFAP immunofluorescence in the spinal cord of WT and TDP43 mice, associated with astrogliosis. Scale bar = 500 µm; detail = 50 µm.

**FIGURE 2.**
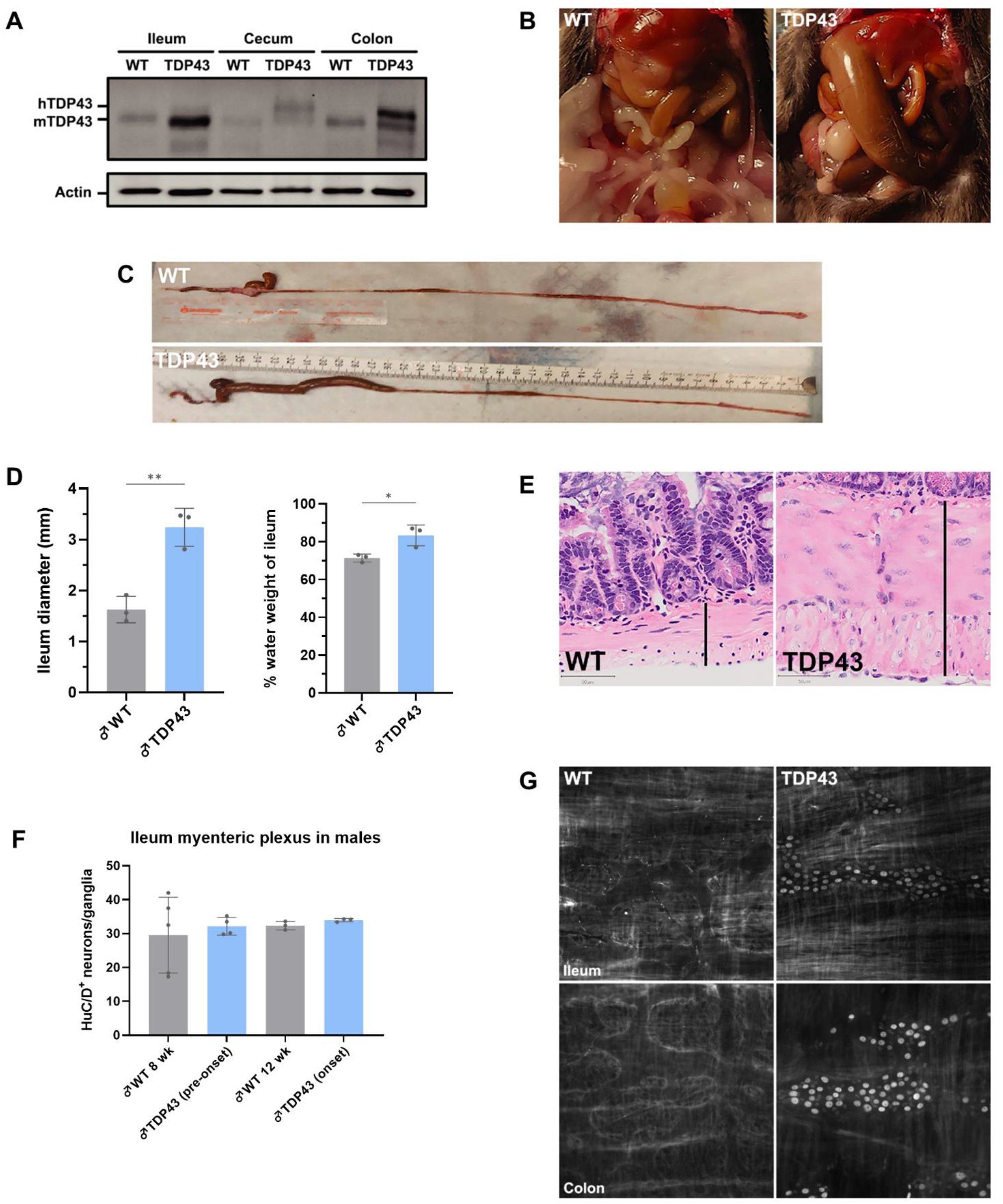
Late-onset alterations in the gastrointestinal tract and enteric system of ALS-linked TDP43 mice. (A) Representative western blot images showing expression of endogenous mouse TDP43 (mTDP43) in separate gut lysates. (B) Representative images of gut morphology in male wild type (WT) and TDP43 mice at 14 weeks of age (end-stage), where the ileum is enlarged. (C) Entire unfolded gut of WT and TDP43 mice. (D) Ileal diameter and percentage of water weight in the terminal ileum in male TDP43 mice at 14 weeks of age (end-stage) and age-matched WT. Data are shown as mean ± SD, n = 3. \**p* < 0.05, \*\**p* < 0.01, unpaired t-test. (E) Representative images showing the thickness of the muscular layer (black line) in the ileum of WT and TDP43 mice. H&E-stained sections (4 μm). Scale bar = 50 μm. (F) Neurons per ganglia in the myenteric plexus of the ileum of WT and TDP43 mice at 8 and 12 weeks of age. Results are shown as mean ± SD, counts of 15 ganglia per animal, n = 3-5, one-way ANOVA test with posthoc Tukey’s test. (G) Immunostaining of human mutant TDP43 protein in the myenteric plexus of ileum and colon of WT and TDP43 mice.

As environmental conditions are critical for the phenotypes of this mouse model ^27–29^, we next analyzed the pathology, motor behavior and survival of our mice. Contrary to previous studies ^27,28^, we consistently observed motoneuron disease (determined by neurological score indicative of hindlimb dysfunction, gait abnormality, kyphosis, and lethargy, in addition to reduced performance in the rotarod assay) in both male and female TDP43 mice. We then focused on male TDP43 mice as they are documented to show severe gut obstruction without motoneuron disease ^27,28^. We performed counts of large motoneurons in the ventral horn of lumbar region of the spinal cord and noted a ∼30% loss of these neurons in male mutant mice (Fig.1C, D), accompanied by astrogliosis in the ventral and dorsal horns, as detected by western blot and immunofluorescence staining, (Fig. 1E, F). These results indicate that CNS histopathology and motoneuron degeneration of ALS-linked TDP43 explain motoneuron dysfunction and disease in our mice.

The ileal enlargement documented in young mice in previous studies ^27,28^ was only detected at the end-stage of the disease in both male and females (Fig. 2B-D). This was accompanied by increased water content in the ileum and thickening of the intestinal wall (Fig. 2D, E). The late appearance of the gut phenotype is consistent with the absence of neuronal death in the myenteric plexus before disease onset (8 weeks of age) and around disease onset (12 weeks of age) in male TDP43 mice (Fig. 2F, G). Thus, in our facility, the TDP43 mice develop ALS-like phenotype, and gut dysfunction is only observed in animals close to end-stage, who succumb with severe neurodegenerative disease.

### Sex differences in the lifespan of TDP43 mice are caused by motoneuron disease

We observed significant sex differences in motoneuron disease progression between male and female TDP43 mice: males develop motoneuron disease faster and their lifespan is shorter than their female counterparts (male TDP43: 101 ± 24 days, n = 89 vs female TDP43: 170 ± 75 days, n = 51; Fig. 3A), with some female animals surviving over 300 days. At 14 weeks of age (around disease onset), male TDP43 mice had worse neurological score, performed poorly on the rotarod assay and showed reduced motoneuron counts in the lumbar spinal cord when compared to age-matched female TDP43 mice, and male and female wild type mice (Fig. 1C, 3B). Female TDP43 had similar number of large spinal motoneurons when compared to age-matched female WT mice. Importantly, TDP43 mice of both sexes reached disease end-stage with all mice in the dataset displaying ALS-like phenotype, and with varying levels of the gut phenotype. Our data differ from previous data showing sex differences in lifespan attributed to the predominant gut obstruction in male TDP43 mice vs the slower motoneuron disease in female TDP43 mice ^27–29^. In our study, all mice were fed with the same food, precluding an effect of diet on the phenotypes and sex differences.

**FIGURE 3.**
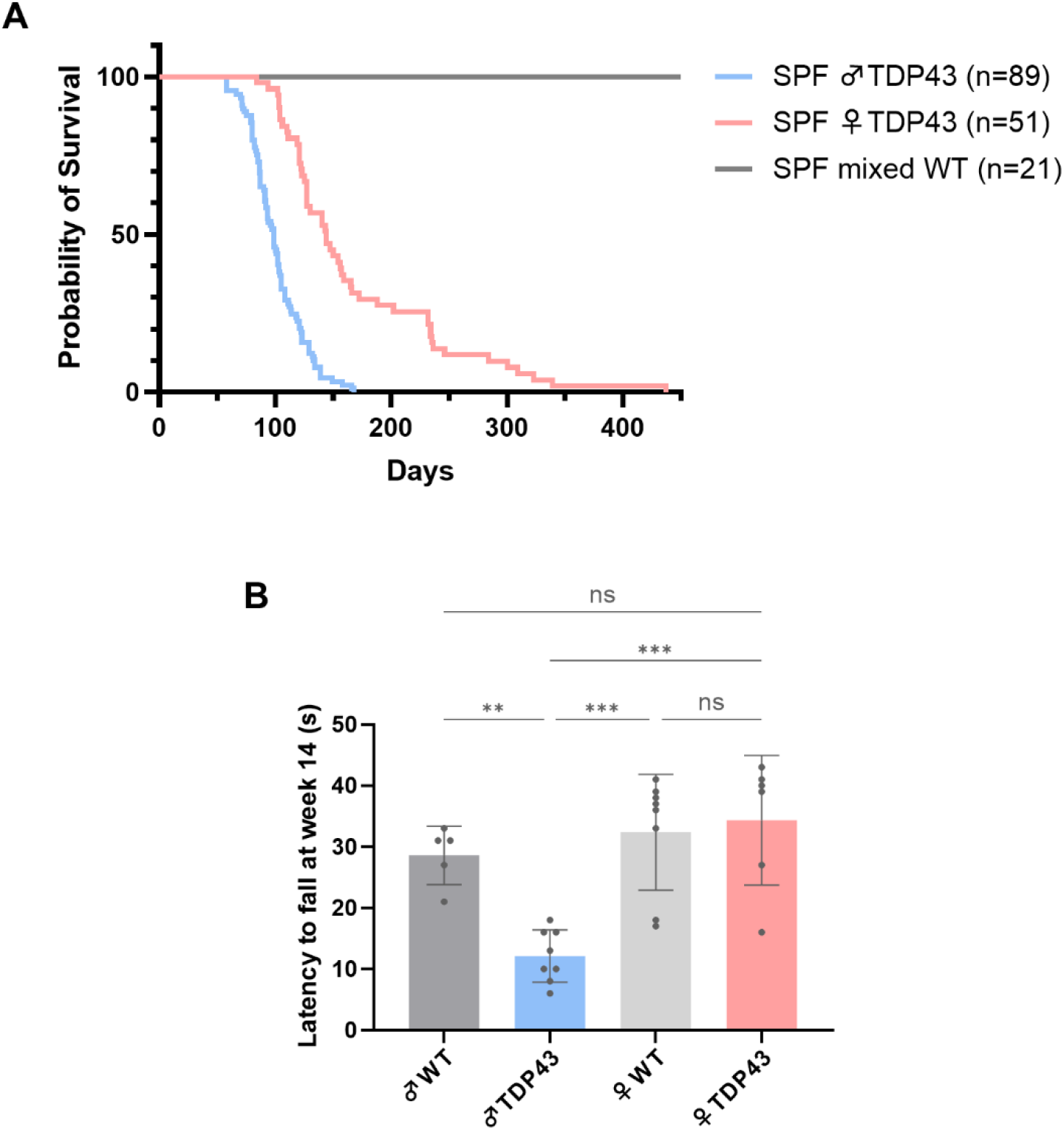
Sexual dimorphism in motoneuron disease onset and lifespan in ALS-linked TDP43 mice. (A) Kaplan-Meier survival graph of SPF TDP43 male and female mice compared to WT mixed-sex mice. Log-rank (Mantel-Cox) test: SPF ♂ TDP43 vs. SPF ♀ TDP43, *p* < 0.0001. (B) Rotarod test at 14 weeks of age. Reduced latency to fall is indicative of impaired motor coordination. Each result is an average of three runs ± SD, n = 5-8, \*\**p* < 0.01, \*\*\**p* < 0.001, one-way ANOVA test with posthoc Tukey’s test.

### The gut microbiota protects against TDP43 toxicity and contributes to sexual dimorphism and pathogenesis in ALS-linked TDP43 (A315T) mice

To determine the role of the gut microbiota in TDP43 mice, we first generated and raised germ-free (GF) TDP43 mice in the International Microbiome Centre at the University of Calgary. These animals were extremely hard to generate with a failure rate of ∼90% at the time of embryo transfer and implantation, probably due to the response trigged by mutant TDP43 in the foster mothers. Over a 36 month period, 9 GF TDP43 mice were produced (5 males and 4 females), and all died precociously (male: 59 ± 3 days, n = 3 vs female: 55 ± 10 days, n = 4 (Fig. 4). We phenotypically and behaviorally characterized the progressive motoneuron disease in seven of the nine GF TDP43 mice. This corresponds to ∼50 days earlier than the male SPF-TDP43 mice and up to 250+ days when compared to longest-lived female SPF-TDP43 mice. We were able to perform the rotarod test on only one GF TDP43 male (58 s to fall) and two GF TDP43 females (70 s to fall), but they performed worse than age-matched WT GF mice (105 s to fall, n = 3). Despite there being few mice, there was no evidence of sexual dimorphism in GF-TDP43 mice (Fig. 4).

**FIGURE 4.**
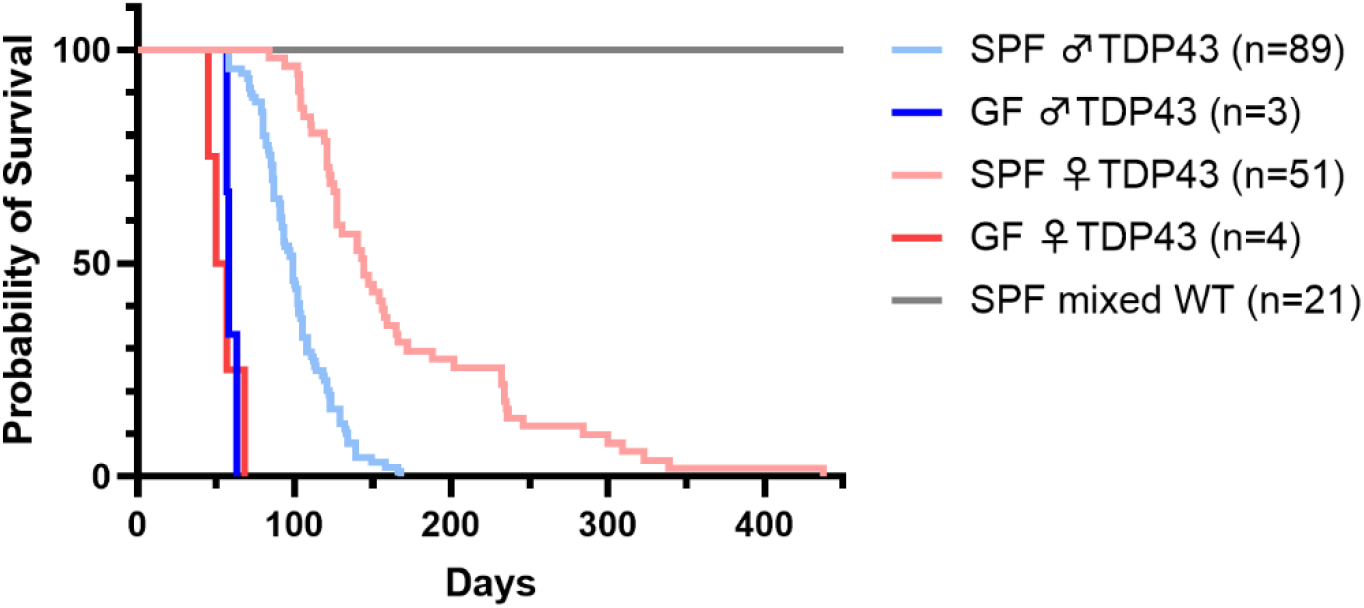
TDP43 mice raised in germ-free conditions have shortened lifespan and show no differences in survival between sexes. Kaplan-Meier survival graph of GF male and female TDP43 mice compared to SPF TDP43 mice. One-way ANOVA test with posthoc Tukey’s test: SPF ♂ TDP43 vs. SPF ♀ TDP43, *p* < 0.0001; SPF ♀ TDP43 vs. GF ♀ TDP43, *p* < 0.0001; GF ♂ TDP43 vs. GF ♀ TDP43, *p* = ns.

To further determine the role of the gut microbiota in TDP43 mice, we next treated animals with a cocktail of antibiotics ((Abx) neomycin, ampicillin, vancomycin and metronidazole) to deplete gut bacteria ^32–34^. Quantification of bacterial DNA in fecal material from mice treated with this cocktail of antibiotics in a pilot study confirmed the depletion of gut bacteria (Fig 5B). Abx-treated 5 weeks old male and female TDP43 mice show much shorter lifespan (male: 49 ± 3 days, n = 10; female: 65 ± 10 days, n = 13) when compared to their mutant counterparts treated with water (male: 96 ± 13 days, n = 13; female: 169 ± 57 days, n = 8 (Fig. 5A)). Weekly rotarod monitoring for motor function detected a significant decline in males as early as one week after the start of the treatment, whereas Abx-treated female TDP43 took a week longer to reach similar decline (Fig. 6A, B). In contrast, both male and female TDP43 treated with water displayed motor deficits much later, with a faster decline for male vs female animals and a more striking difference between sexes. Counts of motoneurons in the ventral horn confirmed the accelerated neurodegeneration in Abx-treated male and female TDP43 mice (male: 32 ± 3 neurons per section; female: 30 ± 5 neurons per section (Fig. 6C)), when compared to age-matched TDP43 mice treated with water (male: 43 ± 6 neurons per section; female: 44 ± 4 neurons per section) and WT mice (male: 41 ± 4 neurons per section; female: 44 ± 4 neurons per section). End-stage TDP43 mice of both sexes were used as positive controls for motoneuron counts (Fig. 6C), thereby confirming the faster neurodegeneration in the Abx-treated mutant mice. Wild type control animals treated with Abx (n = 3-4 per sex) did not show motoneuron disease, indicating that Abx treatment did not cause the ALS phenotype *per se*. Importantly, the striking difference in average survival between male and female TDP43 mice was almost completely abolished with the Abx treatment, suggesting a key role for gut bacteria in sex differences in survival. Thus, using the complementary approaches of germ-free mice and antibiotic treatment to deplete the gut microbiota we discovered that 1) the gut microbiota is protective against TDP43 toxicity and 2) contributes to sexual dimorphism and pathogenesis of ALS-linked TDP43 mice.

**FIGURE 5.**
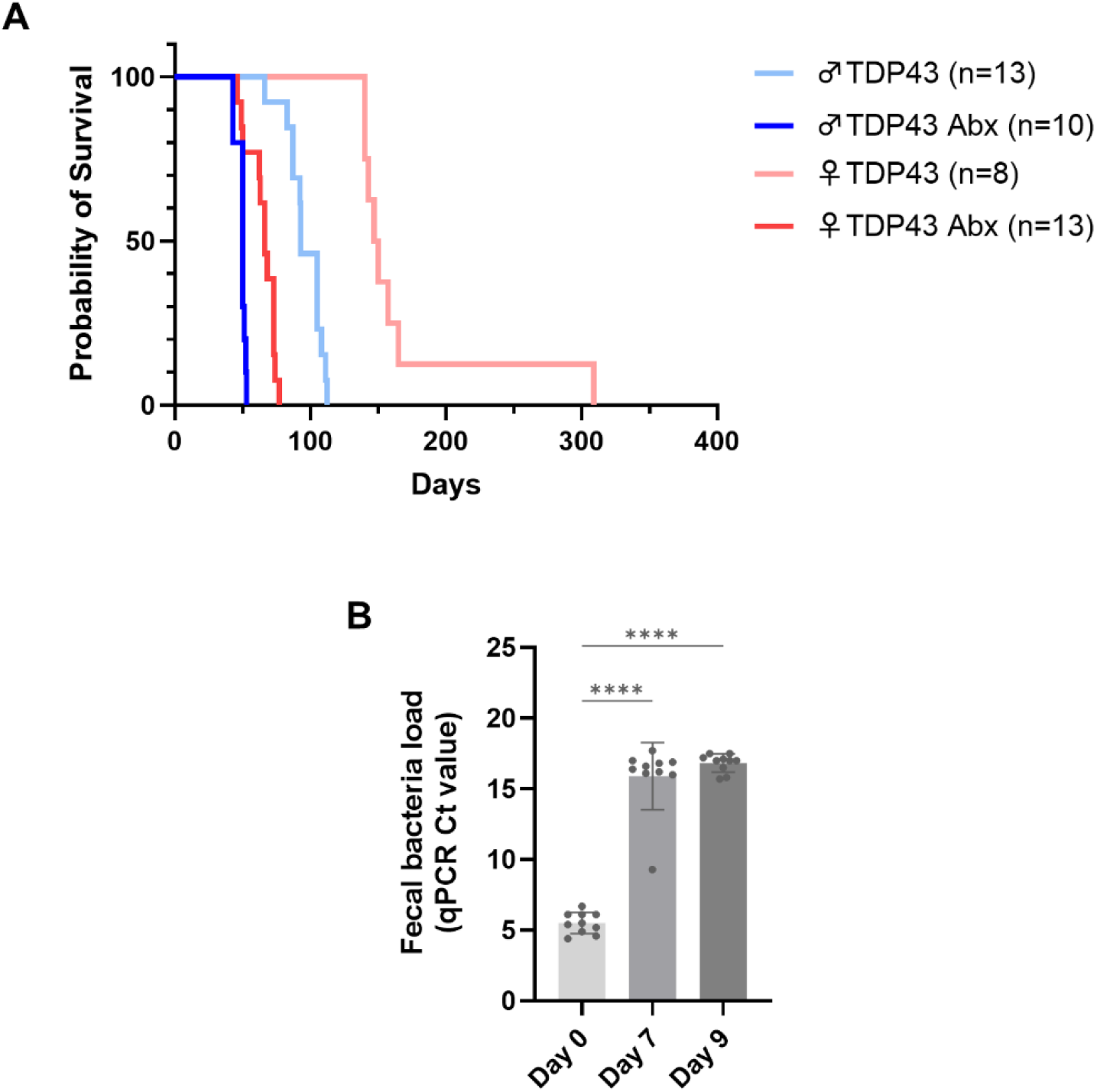
Depletion of gut microbiota with Abx treatment shortens the lifespan of TDP43 mice and abrogates sex differences in survival. (A) Kaplan-Meier survival graph of Abx-treated TDP43 male and female mice compared toTDP43 mice on water. One-way ANOVA test with posthoc Tukey’s test: ♂ TDP43 vs. ♀ TDP43, *p* < 0.0001; ♂ TDP43 vs. ♂ TDP43 Abx, *p* = 0.0006; ♀ TDP43 vs. ♀ TDP43 Abx, *p* < 0.0001; ♂ TDP43 Abx vs. ♀ TDP43 Abx, *p* = ns. (B) Fecal bacteria load of mixed-sex WT mice treated with Abx, showing the effectiveness of the Abx protocol. Data presented as qPCR Ct values. Mean ± SD, n = 10, \*\*\*\**p* < 0.0001, one-way ANOVA test with posthoc Tukey’s test.

**FIGURE 6.**
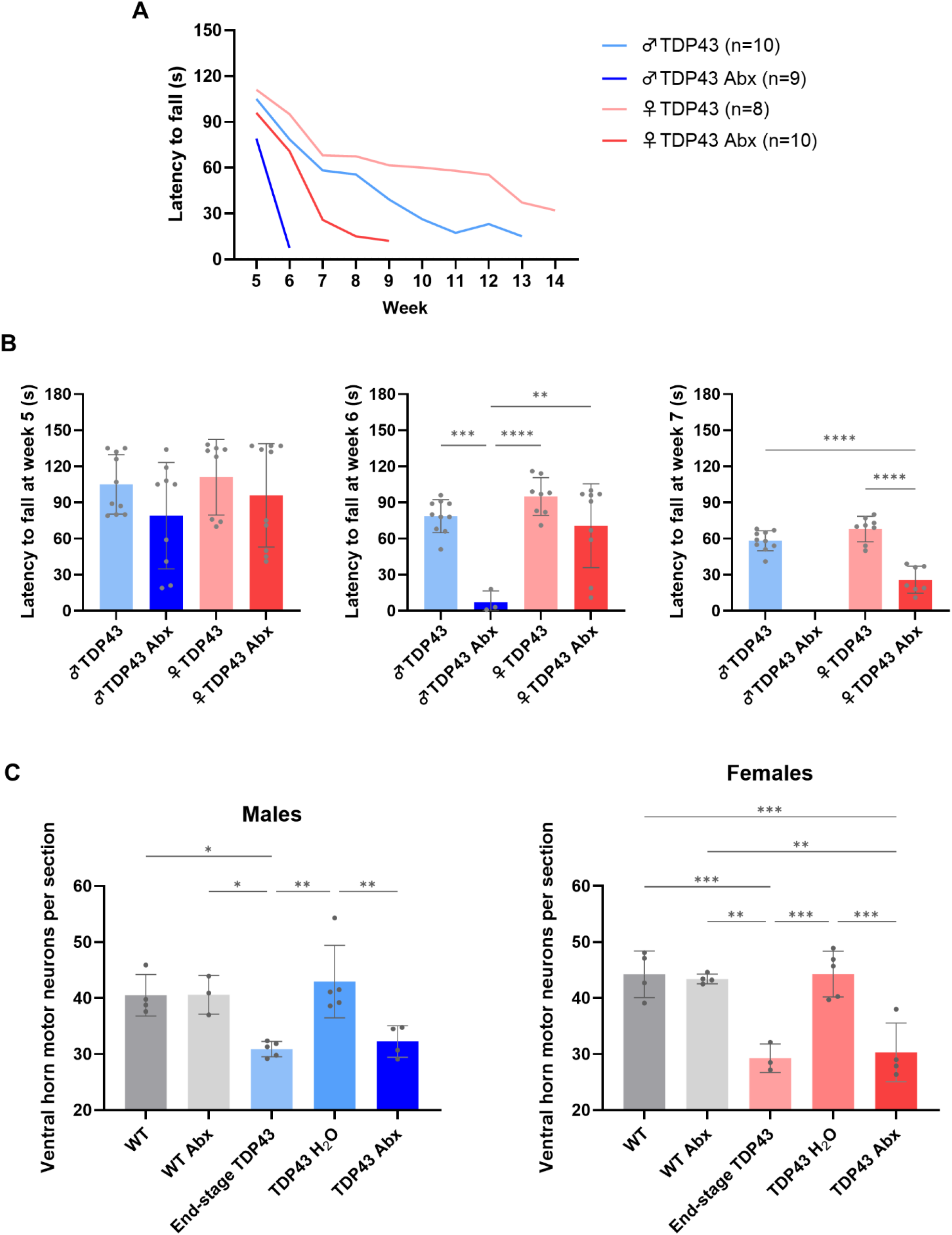
Depletion of gut microbiota with Abx treatment accelerates motoneuron disease and neurodegeneration in the lumbar spinal cord. (A) Weekly progression of rotarod test performance in TDP43 male and female treated with water or Abx. Reduced latency to fall is indicative of impaired motor coordination. (B) Detailed results of rotarod progression at weeks 5, 6 and 7. Each data point is an average of three runs ± SD, n = 8-10, \*\**p* < 0.01, \*\*\**p* < 0.001, \*\*\*\**p* < 0.0001, one-way ANOVA test with posthoc Tukey’s test. (C) Nissl-stained motoneuron count in the ventral horn of the lumbar spinal cord of TDP43 male and female mice treated with Abx. The WT, WT Abx and TDP43 H2O groups are age-matched to the TDP43 Abx mice of the corresponding sex; the end-stage TDP43 and TDP43 Abx groups are end-stage mice (see survival curves in Fig. 5). The number of motoneurons was obtained from counts of 20 sections (16 μm) per animal. Results are shown as mean ± SD, n = 3-5, \**p* < 0.05, \*\**p* < 0.01, \*\*\**p* < 0.001, one-way ANOVA test with posthoc Tukey’s test.

### Castration did not attenuate motoneuron disease in male TDP43 mice

Sex hormones are commonly viewed as drivers of sexual dimorphism and their interactions with the gut microbiota are described (for a review, see ^9^). Male sex hormones (i.e., androgens) produced by the gonads have been hypothesized to play a dual role in ALS, predominantly detrimental (reviewed in ^9^). Through body-brain interactions, these hormones are believed to affect both peripheral tissues (e.g. muscles), systems (e.g. immune) and the CNS ^9^, thereby modulating disease manifestations. To determine whether gonadal-derived androgens contribute to pathogenesis and sexual dimorphism in male TDP43, we performed castration in 5 weeks old mice and monitored them for motoneuron disease progression and lifespan. We found that castration did not alter the course of motoneuron disease in TDP43 mice (castrated: 99 ± 11 days, n = 8 vs non-castrated: 102 ± 9 days, n = 9; non-significant (Fig. 7)). These results indicate that male gonadal-derived sex hormones do not participate in disease pathogenesis in this model.

**FIGURE 7.**
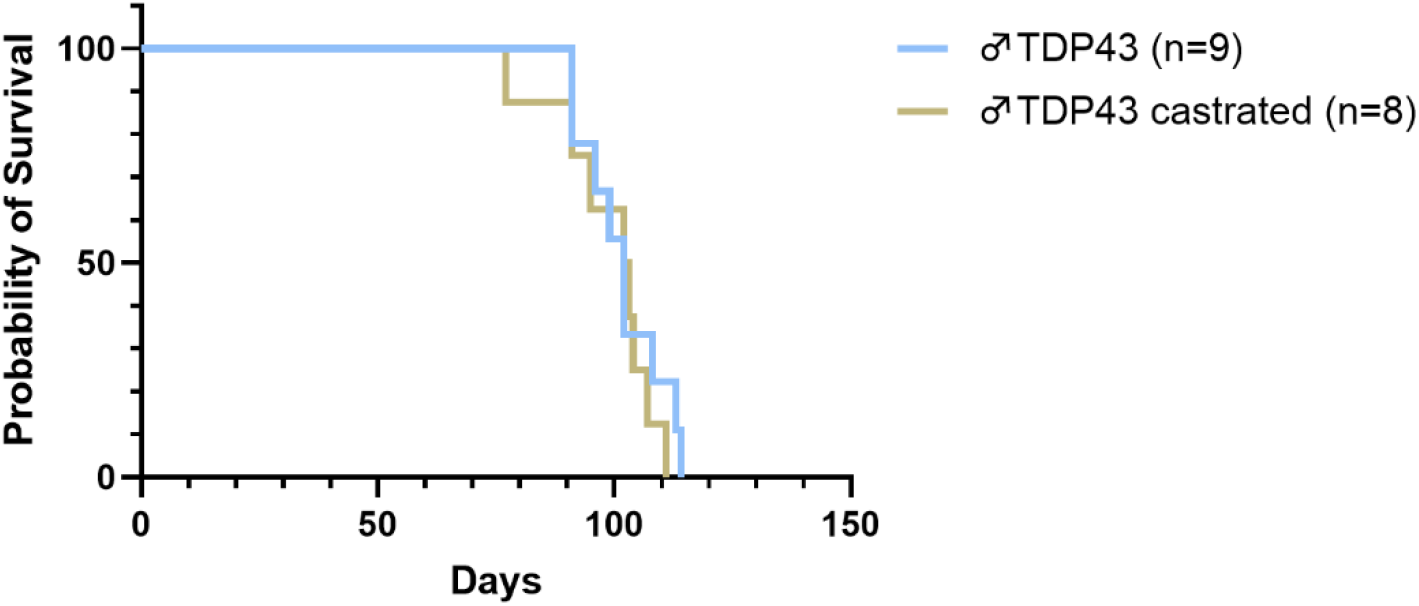
Removal of gonadal male sex hormones by castration does not change disease pathology in TDP43 mice. Kaplan-Meier survival graph of male TDP43 mice castrated at 5 weeks of age compared to non-castrated TDP43mice. Log-rank (Mantel-Cox) test: ♂ TDP43 vs. ♂ TDP43 castrated, *p* = ns.

## Discussion

Several mechanisms have been proposed to explain the sexual dimorphism in ALS including hormonal differences, variability in the blood brain barrier permeability, glial mechanisms and the distinct ability to mount an efficient immune response (for a review, see ^9^). Our study suggests that the gut microbiota is also a key contributor to the sexual dimorphism observed in ALS. In humans, microbiota composition varies significantly between males and females ^25,26^, and therefore, such variation may impact the well-described male predominance of the disease ^2,8,9^.

Many laboratories have reported a gut disease in TDP43 mouse lines that occurs prior to the manifestation of motoneuron disease ^27–29^. This limitation has precluded the study of the TDP43 mice in the context of the microbiota. Furthermore, the reported sex differences in lifespan between male and female TDP43 mice was linked to the more severe gut disease in male vs female mice, and not to the ALS phenotype ^27–29^. A recent study by Zhang indicated that probiotics maintain gut and blood-brain barriers, thereby increasing the motor performance of the TDP43 mice ^20^. However, lifespan as well as sexual dimorphism were not investigated in this study ^20^, probably due to the early gut obstruction phenotype.

Our study suggests that environment is a key determinant in the expression of the motoneuron disease in both male and female TDP43 mice. In our SPF facilities, male and female TDP43 mice develop motoneuron disease prior to gut obstruction and ileal enlargement but at different rates, with males progressing much faster. The loss of gut microbiota not only precipitates disease onset and shortens lifespan, it also abolishes the sex differences in these motoneuron degeneration phenotypes. Our findings are reminiscent of the protective role of gut bacteria in the mutant SOD1 mice ^14^, with the important difference that sexual dimorphism is not found in this mouse line (unpublished observations from the Elinav lab).

How the microbiota confers protection against TDP43 toxicity remains to be investigated. One possibility is through interactions with female sex hormones. There are bidirectional interactions between the gut microbiota and the hormonal systems. For instance, the fecal levels of oestradiol and progesterone in germ-free mice are lower than in SPF mice, suggesting an involvement of gut microbiota in regulation of sex hormones ^35^. On the other hand, gut microbiota is a producer of β-glucuronidase, an enzyme that can reactivate metabolized estrogen by reversing the glucuronidation process, and therefore increases the number of free estrogens in the body ^36^. Interestingly, castration had no effect on the motoneuron disease onset and lifespan of male TDP43 mice, precluding a role for male gonadal-derived sex hormones in disease expression. Future studies are required to determine how female sex hormones interact with the gut microbiome to perhaps impact disease progression in TDP43 mice. Ovariectomy experiments in female TDP43 mice should provide some answers to these questions.

In sum, we propose that the composition of the gut microbiota represents a novel environmental and sex-dependent factor that modulates the pathogenesis of ALS-linked TDP43 mice. As the gut microbiota differs significantly between males and females, it may represent a biological entity to express the sex differences in ALS. Sequencing the gut microbiota of male and female TDP43 mice will provide a better understanding of the role of specific gut bacteria in this mouse model. Ultimately, the work in TDP43 mice will shed new light on the gut microbiome-brain mechanisms regulating sexual dimorphism in ALS.

## Materials and Methods

### TDP43 A315T mouse line

Prp-TDP43^A315T^ transgenic mouse model on the C57Bl/6J background (named herein TDP43 mice) originally generated by Baloh and colleagues ^29^ were obtained from Jackson Laboratories (strain #010700). All mice were kept on a 12h light/dark cycle and allowed *ad libitum* access to food and water. They were genotyped by polymerase chain reaction (PCR) prior to experimentation. Genotypes were confirmed by PCR and western blots using antibodies recognizing the human mutated form. For all experiments, 2-4 cohorts of TDP43 and WT littermates of both sexes were used. All mice were housed and handled according to the Canadian Council on Animal Care guidelines and experimentation and approved by the Health Sciences Animal Care Committee at the University of Calgary (protocol AC19-0089). As per the animal care guidelines, end-stage was recorded within 24 h following the inability to right itself within 30 seconds when laid on its back or displayed lethargy, and/or reduced body weight of >20% from initial adult weight (10 weeks old), and lifespan was recorded as such.

### Derivation of germ-free TDP 43 mice

For the GF experiments, Prp-TDP43^A315T^ transgenic mice were re-derived to germ-free status via two-cell embryo transfer. Mice were bred and maintained at the International Microbiome Centre (IMC), University of Calgary, Canada. GF status was routinely monitored by culture-dependent and - independent methods.

### Antibiotic treatment

Antibiotic (Abx) treatment was administered according to previously published protocols ^32–34^. Broad-spectrum Abx mix was diluted in sterilized water and consisted of ampicillin (1 g/L; Sigma-Aldrich, A9518), neomycin (1 g/L; Sigma-Aldrich, N1876), vancomycin (0.5 g/L; Sigma-Aldrich, 94747), and metronidazole (1 g/L; Sigma-Aldrich, M3761). Because of the body weight loss observed by our lab and others when treating mice with metronidazole, we opted to gradually introduce it in the Abx solution as previously described ^32–34^, with the full concentration added on day 10. All the other antibiotics were introduced at full concentration on day 0. Body weight was assessed three times a week to monitor body weight changes, and mice had access to the Abx solution for the duration of the study.

### Castration

Castration was performed in male TDP43 mice at 5 weeks of age. Briefly, mice were injected subcutaneously with Buprenorphine (0.05 mg/kg) and Meloxicam (5 mg/kg), then anaesthetized with isoflurane (induced with 5% and maintained at 2%) throughout the procedure. Surgery was performed by an incision in the abdomen, cauterization of the vas deferens and blood vessels, removal of both testicles and suture of the body wall with wound clips. For the next two days after surgery, mice were given Buprenorphine (0.05 mg/kg) every 12 hours and Meloxicam (5 mg/kg) every 24 hours, subcutaneously. Recovery was monitored for 10 days, after which wound clips were removed. Mouse health and disease progression were then assessed until the experimental endpoint.

### Assessment of motor function and survival

#### Tail suspension

We assessed disease progression with the tail suspension test and assigned a neurological score using established criteria ^37^. Mice were held above their cage by their tail and limb position and movement was observed for 5 seconds. Symptomatic mice displayed collapsed limbs (limbs pointing inwards) and one or both hindlimb clasped, while asymptomatic mice displayed outstretched limbs and a wider range of fast limb movements (Table 1).

**TABLE 1.** Summary of observations used to assess disease progression.

| **Neurological score** | **Description** |
| --- | --- |
| 0 (pre-symptomatic) | Normal; full extension of hindlimbs away from lateral midline when the mouse is held by its tail |
| 1 (first symptoms) | One hindlimb collapsed towards the midline or trembling; normal gait; no weight loss |
| 2 (disease onset) | Both hindlimbs collapsed and clasped; gait abnormality; starts losing weight |
| 3 (paralysis) | Complete loss of extension reflex in both hindlimbs; dragging hindlimbs while walking and less active overall; kyphosis; loss of <20% body weight |
| 4 (humane endpoint) | Mouse cannot right itself when laid on its back within 30 seconds and/or display lethargy; loss of >20% body weight |

Adapted from Hatzipetros et al. <sup>37</sup>

#### Rotarod

Mice were run on an accelerating rotarod, and latency to fall was recorded. The final result was an average of three trials. Results from 14 week mice were obtained using a single-day protocol ^38^ where mice were pre-trained for three trials before the formal test, and during the test, the rotarod accelerated from 4 to 40 rpm over 300 seconds. When possible according to the experiments, routine rotarod tests were performed to obtain more stable performances, following a different protocol (Standford Behavioral and Functional Neuroscience Laboratory): mice were trained for five consecutive days on the rotarod and subsequently tested weekly starting at five weeks of age. During the training phase, mice were run on the rotarod at 5 rpm for 60 seconds for three rounds per session with 10 min intervals. During the testing phase, the rotarod was accelerated from 4 to 40 rpm over 300 seconds. Each mouse underwent three trials per session with 15 min intervals.

#### Survival

Mouse health and disease progression were assessed daily. The end-stage of disease (humane endpoint of the experiment) was defined as when a neurological score of 4 was reached, which is when the mouse could not right itself when laid on its back and/or exhibits lethargy and/or displays weight loss higher than 20% of initial adult body weight (Table 1). At this point mice were euthanized. Based on these endpoints, the Kaplan-Meier survival curve was generated and analyzed using a log-rank test (for 2 groups of mice) or one-way ANOVA (for multiple groups of mice).

### Western blot

Mice were anesthetized and perfused with perfusion solution containing 1 M sodium fluoride, 0.15 M sodium pyrophosphate, and cOmplete^TM^ protease inhibitor cocktail (diluted 1 in 20; Roche, #11697498001) in PBS. Spinal cord, brain, and gut (*i.e.,* ileum, cecum, and colon) tissues were obtained from mutant TDP43 and age-matched WT mice. Tissue samples were lysed on ice using Tris-EDTA SDS lysis buffer and homogenized using a tissue homogenizer and sonicated for 10 seconds (1 s on/1 s off). The samples were centrifuged at 13000 rpm at 4 °C for 15 min and the supernatants were collected. The samples were quantified using the DC protein assay kit (Bio-Rad laboratories, #500-0116). 40 µg of each sample was prepared with a 5x Laemmli sample buffer and heated for 5 min at 95 °C . Samples were loaded onto an SDS-PAGE gel and separated under 130 volt for 1 h on ice. After gel electrophoresis, the gel was placed under 110 volt current for 1 h on ice for transfer to a polyvinylidene fluoride membrane. The membrane was blocked in 5% skim milk for 1 h and probed with a primary antibody overnight at 4 °C (Table 2). The membrane was washed with phosphate-buffered saline solution (PBS) with Triton X-100 (PBST) and probed with Horseradish peroxidase (HRP)-conjugated secondary antibodies, diluted in 5% skim milk in PBST for 1 h at room temperature. The membrane was then washed for a total of 1 h (4 x 15 min each) with PBST. Blots were visualized using ECL chemiluminescence (PerkinElmer, #22360215) substrate and imaged on a ChemiDoc^TM^ imaging system (Bio-Rad Laboratories). Band intensity was quantified using the Image Lab software (Bio-Rad Laboratories) and normalized to loading control (actin or GAPDH).

**TABLE 2.** Antibodies.

| **Antibody** | **Method** | **Dilution** | **Company** | **Catalog #** | **Host** |
| --- | --- | --- | --- | --- | --- |
| TARDBP (TDP43)<br>Mouse | WB | 1:500 | Proteintech | 12892-1-AP | Rabbit |
| TARDBP (TDP43)<br>Human | IHC brain<br>IHC gut | 1:200<br>1:1000 | Abnova | H00023435-M01 | Mouse |
| GFAP | WB<br>IHC | 1:1000<br>1:400 | Chemicon | MAB360 | Mouse |
| Actin | WB | 1:1000 | Chemicon | MAB1501 | Mouse |
| GAPDH | WB | 1:1000 | Abcam | ab9484 | Mouse |
| MAP2 | IHC | 1:200 | Millipore | AB5622 | Rabbit |
| HuC/D | IHC | 1:200 | Invitrogen | A21271 | Mouse |
| Anti-mouse Alexa<br>Fluor 488 | IHC | 1:200 | Jackson<br>ImmunoResearch | 715-546-151 | Donkey |
| Anti-rabbit Cy3 | IHC | 1:100 | Jackson<br>ImmunoResearch | 711-165-152 | Donkey |
| HRP-conjugated anti-mouse IgG | WB | 1:500 | EMD Millipore Corp | AC111P | Sheep |
| HRP-conjugated anti-rabbit IgG | WB | 1:500 | Cytiva | NA934-1ML | Donkey |

WB: western blot; IHC: immunohistochemistry

### Immunohistochemistry

#### Brain and spinal cord

Mice were anesthetized and perfused with 4% PFA solution diluted in PBS. Whole brain and lumbar spinal cord were collected. Tissues were fixed in 4% paraformaldehyde (PFA) (ThermoFisher Scientific) and cryopreserved in 30% sucrose. The cryopreserved tissues were mounted in OCT and sectioned at 10-30 μm slices using a cryostat. Sections were blocked with 5% skim milk in PBST incubated in primary antibody overnight at 4 °C. Slides were incubated in conjugated secondary antibodies and stained with DAPI. Images were obtained using either fluorescence microscope (Olympus VS110 Slides Scanner, Olympus Lifescience) or a confocal microscope (FLUOVIEW FV3000, Olympus Lifescience). Antibodies that were used are described in Table 2.

#### Myenteric plexus

Male TDP43 mice and age-matched littermate WT mice were anaesthetized with isoflurane and euthanized by cervical dislocation. A 3 cm-length piece of distal ileum and proximal colon were collected and dissected to prepare whole mounts of myenteric plexus. First, the intestinal samples were bathed for 10 min in a solution containing 1 μM nifedipine (Sigma-Aldrich) diluted in PBS. Tissues were then opened along the mesenteric border and pinned out in a Sylgard-coated petri dish with the serosal side facing down. Samples were fixed with 4% PFA solution for 24 h at 4 °C. Tissues were then washed with PBS containing 5% sodium azide (3 x 10 min) and stored at 4 °C until further processing. To prepare whole mounts of the myenteric plexus, samples were dissected by stripping off the mucosa/submucosa and the circular muscle layer, leaving a preparation consisting of the longitudinal muscle and associated myenteric plexus. Myenteric preparations were then processed for immunohistochemistry. First, tissues were washed (3 x 5 min) in PBS containing 0.1% Triton X-100 (Sigma-Aldrich) and incubated with mouse anti-TARDBP primary antibody or mouse anti-HuC/D (for total neurons) ^34^ for 48 h at 4 °C. Next, the samples were washed (3 x 5 min) with PBS and incubated with secondary antibody (donkey anti-mouse Alexa Fluor 488) for 1–2 h at room temperature. This was followed by washing (3 x 5 min) with PBS. Whole-mount preparations were mounted with bicarbonate-buffered glycerol on microscope slides and stored at 4 °C under dark conditions for further analyses. All the antibodies used were diluted in a PBS solution containing 0.1% Triton X-100, 0.1% bovine serum albumin, 0.05% sodium azide, and 0.04% EDTA. Tissues were microscopically analyzed using a Zeiss Axioplan fluorescence microscope (Zeiss Canada) and images were captured with a QImaging Retiga-2000R digital monochrome camera (Teledyne Photometrics). Detailed information on the antibodies used can be found in Table 2.

## Morphological and cellular analysis

### Motoneuron quantification

Nissl-stained large motoneurons (diameter >25 µm) in the ventral horn of the lumbar spinal cord region (L3-5) of WT and TDP43 mice at end-stage (14 weeks of age, or when the motor deficits were evident) were counted using the QuPath software. The mean number of motoneurons was obtained from 20 sections per animal with 3-5 animals per genotype. Serial 16 μm sectioning was performed, with every 3rd section placed on a slide, to avoid duplicate counts. Images were taken using the Olympus VS110 Slides Scanner microscope (Olympus Lifescience).

### Hematoxylin & Eosin (H&E) staining of ileum

We collected 2 cm distal ileum from mice to process for H&E staining and examine the morphology of the gut. After fixing the samples in 10% formalin, the samples were sent to Alberta Laboratory Services for paraffin embedding, sectioned into 4 μm-thick sections, and H&E stained. Images of the sections were obtained using the Olympus VS110 Slide Scanner (Olympus Lifescience).

### Gut edema

We obtained terminal ileal tissue from TDP43 mice at end-stage and age-matched WT and measured the weight of the tissue from the mice after euthanasia (wet weight) and after drying the tissue overnight at room temperature (dry weight). We measured the percentage of the weight that is due to water using the following formula: % Water weight = 100 * (wet weight - dry weight)/wet weight.

#### Statistical analysis

Statistical analyses were conducted using GraphPad Prism (GraphPad, San Diego, CA). The survival of mice was calculated using the Kaplan–Meier method, and statistical analysis was performed using a log-rank test or a one-way ANOVA test. In the characterization of the TDP43 mice, an unpaired student’s t-test was applied for comparing two groups, and a one-way analysis of variance (ANOVA) with Tukey’s post hoc analysis was employed to analyze results from more than two groups. For all studies, *p* < 0.05 was considered significant.

## Acknowledgements

The authors thank Cameron Fielding (Clara Christie Centre for Mouse Genomics, University of Calgary) for performing the castration surgery and Laurie Wallace for the support with bacterial quantification after antibiotic treatment. We also thank the Advanced Microscopy Platform (Hotchkiss Brain Institute, University of Calgary) for the support with microscopy and imaging, and the CSM Optogenetics Facility (Hotchkiss Brain Institute, University of Calgary) for the support with the rotarod performance test.

## Author contributions

EJB, SL, KDM, JB, GP, EE, KAS and MDN designed the studies; EJB, SL, YJ, IMR, RP, CMK, MS,

MB and OB conducted experiments and performed data analyses; EJB, SL, KAS and MDN drafted the manuscript. All authors had access to the study data, critically reviewed manuscripts drafts and approved the final manuscript for submission. KAS and MDN obtained funding for the study. MDN provided study supervision and oversight.

